# Fine-scale Inference of Ancestry Segments without Prior Knowledge of Admixing Groups

**DOI:** 10.1101/376137

**Authors:** Michael Salter-Townshend, Simon Myers

## Abstract

We present an algorithm for inferring ancestry segments and characterizing admixture events, which involve an arbitrary number of genetically differentiated groups coming together. This allows inference of the demographic history of the species, properties of admixing groups, identification of signatures of natural selection, and may aid disease gene mapping. The algorithm employs nested hidden Markov models to obtain local ancestry estimation along the genome for each admixed individual. In a range of simulations, the accuracy of these estimates equals or exceeds leading existing methods that return local ancestry. Moreover, and unlike these approaches, we do not require any prior knowledge of the relationship between sub-groups of donor reference haplotypes and the unseen mixing ancestral populations. Instead, our approach infers these in terms of conditional “copying probabilities”. In application to the Human Genome Diversity Panel we corroborate many previously inferred admixture events (e.g. an ancient admixture event in the Kalash). We further identify novel events such as complex 4-way admixture in San-Khomani individuals, and show that Eastern European populations possess 1 – 5% ancestry from a group resembling modern-day central Asians. We also identify evidence of recent natural selection favouring sub-Saharan ancestry at the HLA region, across North African individuals. We make available an `R` and *C*++ software library, which we term MOSAIC (which stands for MOSAIC Organises Segments of Ancestry In Chromosomes).

## Admixtur

Occurs when reproductive isolation between groups allows genetic divergence via genetic drift and random mutation, followed by mixing of the diverged groups to form new populations. Such genetic admixture is near ubiquitous in observed human populations (Hellenthal *et al*. 2014; Loh *et al*. 2013; Patterson *et al*. 2012) and indeed other species including cattle (Upadhyay et al. 2016), bison (Musani *et al*. 2006), and wolves (Pickrell and Pritchard 2012).

Genome-wide summaries can reveal not only complex relationships between modern populations but also details of their demographic histories (Pickrell and Pritchard 2012; Hellenthal *et al*. 2014; Peter 2016) while accurate inference of local ancestry can be used to correct for population structure in association testing (Diao and Chen 2012; Xu and Guan 2014), detect selection (Zhou *et al*. 2016), and can be used for mapping disease loci (Zhang and Stram 2014).

Due to the process of recombination, contiguous chunks of admixed individuals’ genomes are inherited intact from one mixing population or another. In the second generation following the initial admixture, chromosomes from distinct ancestral groups begin to recombine, and so the expected length of these chunks (in units of Morgans) will be one (by definition) and (neglecting crossover interference) chunk lengths can be modelled using an exponential distribution with rate parameter 1. In each subsequent generation, recombination further breaks down these chunks so that the chunk lengths (if they could be observed) are distributed according to an exponential distribution with rate parameter one less than the number of generations since admixture.

To fully characterize admixture for the above purposes, we need to infer: (1) Whether a group of individuals are admixed (2) The component / mixing groups (3) The timing of the admixture event(s) (4) Which segments of the admixed genome are inherited from each mixing group. Typically we lack prior knowledge of each of these points and we do not have access to representative samples of the mixing groups, as these are often no longer present (without drift) in modern samples.

A wide variety of approaches to model admixture have been developed in recent years. STRUCTURE (Pritchard *et al*. 2000) clusters similar genomes together by fitting a mixture model using Gibbs sampling and STRUCTURE 2.0 (Falush *et al*. 2003) extended this model to allow for admixed individuals using a Hidden Markov Model (HMM) that allowed for linkage along the genome. A drawback of these and similar Sohn *et al*. (2012) approaches is that they do not attempt to model linkage disequilibrium (LD), because SNPs within each source population are assumed to be independent, meaning they are not maximally powerful for inferring ancestry segments, particularly for subtle admixture events.

Other approaches focus on dating/characterizing admixture events, without performing local ancestry estimation. In the ALDER model (Loh *et al*. 2013) the exponential decay of ancestry segments is estimated as a function of genetic distance, allowing dating of admixture events. GLOBETROTTER (Hellenthal *et al*. 2014) uses a related approach for dating events by leveraging haplotype data, accounting for LD, but also infers admixture proportions and properties of the ancestral mixing groups, by quantifying their relationships with modern observed populations, and can handle multi-way admixture. In common with other approaches (some discussed below), GLOBETROTTER incorporates LD between nearby SNPs by fitting a haplotype copying model (Lawson *et al*. 2012) closely related to the Hidden Markov model introduced by Li and Stephens (Li and Stephens 2003). Here, “target” chromosomes of interest are formed as a mosaic whereby they imperfectly “copy” segments of DNA from donor haplotypes, according to a HMM. See Gravel (2012) for a review of this and other local ancestry models. The subsequent copying profiles (both global and locally along the genome) are analysed and decomposed. Admixture times are then inferred by fitting curves measuring the correlation in copying along the genome: the relative probability of copying from pairs of donor populations is estimated at increasing genetic distances.

Finally, several statistical algorithms (e.g. (Churchhouse and Marchini 2013), (Guan 2014) (Price *et al*. 2009)), have been developed to identify local ancestry segments while accounting for LD: however, these rely on pre-specification of the number of mixing groups, and the inclusion of groups of donor samples identical to, or at least closely related, to these mixing groups. Of these, HapMix (Price *et al*. 2009) models both pre- and postadmixture recombination via a two-layer HMM, using an algorithm related to that we develop here. However, HAPMIX only allows for two admixing groups and the user must supply known surrogates for both ancestries (a low mis-copying rate is allowed for in the model).

### MOSAIC overview

Here, we introduce an approach that unlike existing approaches is able to identify local ancestry segments, without requiring prior knowledge of mixing groups. Instead, the details of admixing populations are inferred as part of the algorithm. Our approach: (1) Takes one or more perhaps admixed genomes (2) Compares to previously labelled (e.g. by region of origin, or genetic clustering) groups/reference panels of additional individuals (3) Identifies and characterises segments of local ancestry for admixture of arbitrary numbers of populations. Note that we do not require that the unobserved admixing ancestral groups are a close match with the observed labelled groups, but rather we learn the genetic relationships between them. We exploit LD information to decompose the genome into segments and use an HMM algorithm, similar in spirit to that of HapMix, which forms a special case. Each admixing population shares haplotypes across all reference panels, copying from each panel according to a set of weight parameters inferred by the method. For example, for data analysed in (Hellenthal *et al*. 2014), we explore admixture within the Hazara individuals (see Hazara 2-way), by comparing their genomes to those of members of the 94 other labelled groups within the dataset (Hellenthal *et al*. 2014). We find Hazara individuals possess admixture segments from two groups, one preferentially copying from donors that are North East Asian, and one from Central/South Asian donors, matching previous findings (e.g. (Hellenthal *et al*. 2014)).

To avoid phasing errors scrambling ancestry switch signals within the inferential algorithm, MOSAIC iteratively updates haplotypic phase, and infers time since admixture via the fitting of exponential decay coancestry curves. We also estimate the drift between the unobserved ancestral groups and other groups, by constructing partial genomes from the admixed individuals themselves, representative of the original non-admixed ancestral individuals. These can then be compared to other populations, allowing estimation of e.g. F_st_.

Our software, MOSAIC, returns parameter estimates, local ancestry estimates along the genome, co-ancestry curves (including the best fit exponential curve), ancestry informed phase of the target haplotypes, and Fst estimates between the ancestral mixing groups and between the mixing groups and each panel.

## Materials and Methods

The inputs are haplotypes from labelled sub-populations (panels), target admixed haplotypes, and recombination rate maps. Typically the phasing of the haplotypes is accomplished algorithmically (here we have used SHAPEIT2 (Delaneau *et al*. 2013)) but we note that phasing errors in the donor haplotypes will have minimal impact on inference and we correct phasing errors in the admixed targets within our inferential algorithm.

### Illustration of the Model

Figure 1 depicts our model in graphical form, with further details below. Figure 1a depicts the phased part of our approach but we correct for phasing errors that scramble the ancestry as described in Re-Phasing and Figure 1b. In the figure, there are 12 phased haplotypes that belong to 4 modern populations (panels). These will act as potential surrogates for the target 2-way admixed haplotype. We do not begin with knowledge of surrogate to admixing source relationships but learn it from the data. The matrix of copying probabilities ***μ*** parameterises this; ***μ***_*pa*_ is the conditional probability that a haplotype from panel *p* is chosen to copy from given the local ancestry is *a*. The columns are constrained to sum to unity and we scale these probabilities when used in our HMM by *N_p_*, the number of donor haplotypes in panel *p*. We infer the values in this matrix directly from the data and thus do not required the user to have any knowledge of the panel to mixing ancestry relationship a-priori. If a panel does not contain useful surrogates for the mixing groups then the corresponding row of this matrix will tend to zero. If there are panels that represent well the mixing groups the corresponding elements will tend towards one. Most usefully, if two or more panels contain good surrogates for an ancestry then both will share the conditional probability mass with obvious post-fit interpretation. Finally, if there is a panel that is similarly admixed to the target individuals this will be reflected as a row in the copying matrix ***μ*** with multiple non-zero entries.

**Figure 1.**
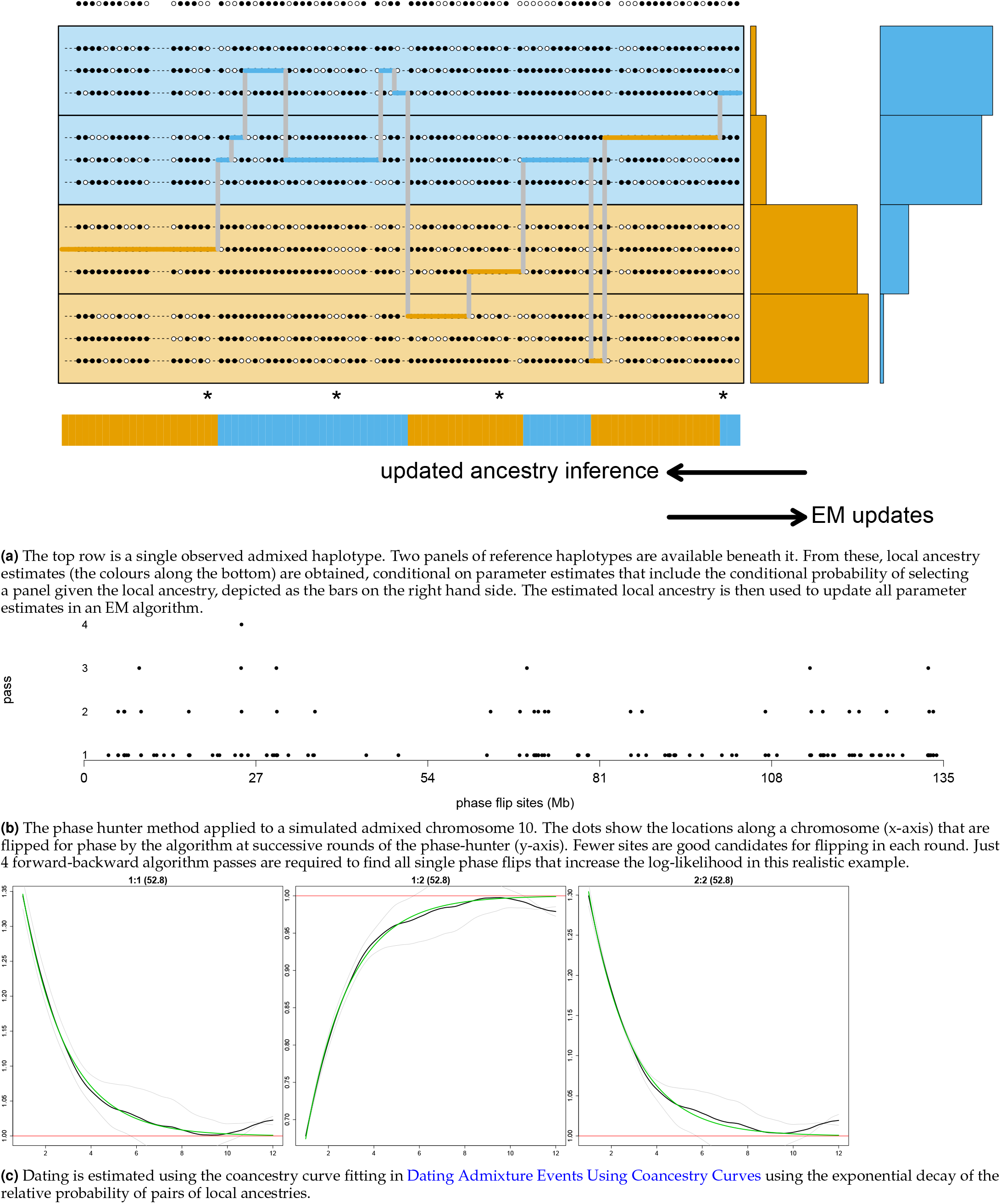
MOSAIC proceeds by rounds of thin (see Thinning), EM (see EM Updates), phasing (see Re-Phasing). 1a is a cartoon version and 1b and 1c depict the realistic simulations used to test the approach in Simulation Studies.

As per all methods derived from Li and Stephens (2003), we model a target haplotype as a mosaic of donor haplotypes in a Hidden Markov Model framework. The hidden states in our model consist of two layers; each locus (or gridpoint; see Gridding on Genetic Distance) has both a hidden local ancestry (blue or red in the Figure) and the other hidden state is which donor haplotype is being copied from. Switches in which ancestry the target is locally in are rare and are parameterised with the rates matrix **Π**.

### Two Layer Hidden Markov Model

Our approach may be viewed as a combination of HapMix (Price *et al*. 2009) and GLOBETROTTER (Hellenthal *et al*. 2014). As per HapMix, admixture is directly incorporated into the HMM. However, unlike HapMix, our model works with any multiple of ancestry sources ^1^ and is more flexible; not only do we allow for a rich variety of dependency between latent ancestral sources and labelled modern populations, but we do not require prespecification of these dependencies (see Transitions for how we parameterize these relationships). As in GLOBETROTTER, MOSAIC infers the relationship between modern populations and ancient unseen mixing populations from the data. The key difference is that our method builds these relationships directly into the HMM that uncovers accurate local ancestry estimates along the genome whereas GLOBETROTTER fits a mixture model to the output of an ancestry unaware HMM. We therefore view our method as the best of both worlds between HapMix and GLOBETROTTER.

#### Gridding on Genetic Distance

In order to perform the EM updates to parameter estimates of the HMM, counts of all possible switch types (ancestry and haplotype copying) need to be enumerated. Given unequal spacings of markers, this necessitates integrating over an infinity of possible switches along the continuous genome. We therefore impose an even grid on recombination distance along each chromosome. This is extremely fine (60 gridpoints per centimorgan) and does therefore not resemble the single-ancestry windows imposed by some existing admixture models.

We can then make the assumption that recombination only happens between successive gridpoints. This will be quite accurate for the time frame on which we focus and greatly simplifies the mathematics of our inference. It is important to note that there can now be 0,1, or multiple SNPs at any given gridpoint and the HMM is defined at all gridpoints. Calculation of the agreement between any donor haplotype and any recipient haplotype is then based upon a one-off calculation of the number of matching SNPs and the number of observed SNPs at each grid-point. Phasing is also calculated only at gridpoints. Crucially, the density of markers will not then greatly impact the speed of our inference, post the reduction of the data to the gridded version. This means that MOSAIC scales sub-linearly with the number of sites so we can analyse sequencing data.

#### Transitions

We jointly model ancestry and haplotype copying chunks along the genome using a two-layer Hidden Markov Model (HHM). The first layer involves ancestry switches along the genome and the second layer switches between copied haplotypes along the genome. Ancestry switches occur at a slower rate than the haplotype switches as they only occur post-admixture and each ancestry switch enforces a haplotype switch in the model. The probability of making a switch from ancestry *b* to ancestry *a* between successive gridpoints is parameterised in our model as 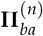 in target individual n. Note that this individual specific *A* × *A* matrix of switch rates encompasses all that the model knows about the admixture event. i.e. the time of the event and the mixing proportions. In Dating Admixture Events Using Coancestry Curves we outline why we rely on coancestry curve fitting to give a final statement about the number of generations since admixture.

The probability of switching to any donor haplotype depends on the size of the panel, and the underlying local ancestry at the gridpoint. We parameterise by ***μ***_*pa*_ the probability of selecting from panel *p* when the local ancestry is *a*. Finally, we denote the recombination within ancestry probability with *ρ*.

The transition probability of making a switch from (ancestry, haplotype) pair (*b, h_q_*) to (*a, h_p_*) where *h_p_* is a donor haplotype *h* in panel *p* for target individual *n*, is given by:

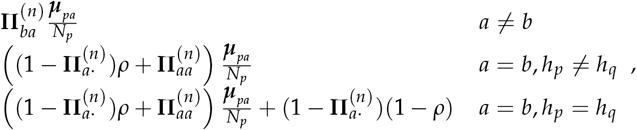

where 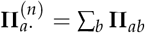.

**Π**^(*n*)^ matrices are specific to the admixed target individuals whereas ***μ*** and *ρ* are assumed to be common to the entire admixed population. All possible choices are all available to our model (and code) and can prove useful; for example when a subset of admixed targets have undergone a markedly different admixture event to the rest they are more usefully analysed with two sets of parameters. The interpretation of the above choice is that the admixture is well characterized as being the mixing of a single set of well defined ancestral populations but each target individual may have experienced the mixing at a different point in their history (and in a different ratio). In this work we assume a single scalar *ρ*; we have experimented with ancestry specific versions of this parameter but the vector ***ρ*** is then confounded with **Π**.

#### Emissions

We deal with biallelic SNP data (denoted with a ***ϒ***) and we use *θ* to parameterize the emission probability of a 1 at gridpoint g when copying donor haplotype *h* as

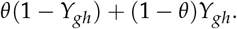

Thus *θ* is the probability of a pointwise discrepancy between the allele of the haplotype being locally copied and the allele of the copying haplotype i.e. the miscopying rate. Note that for notational simplicity we have suppressed here the index of the panel from which that haplotype comes. The panel being copied does not impact our calculation; we could allow this to account for genetic drift between the ancestral groups and the modern reference panels, however this would be confounded with the copying probabilities ***μ***.

As we have moved the observed markers to a grid, each gridpoint may have zero, one, or multiple emissions (observations). This is simply handled by assuming a product of emission probabilities for multiple observations (or a sum over the log-probabilities in practice). For gridpoints with no observations, the emission probability is simply 1. This has the additional benefit of allowing our model to handle missing data seamlessly; there is simply a lower count of observations at a gridpoint when donor SNPs are missing.

### Algorithm

Our inferential algorithm comprises initialisation of all parameters (see Appendix Initialisation), followed by a loop over successive rounds of thinning (see Thinning), re-phasing (see Re-Phasing), and EM updates of the parameters (see EM Updates). We find that a low number (5 for results presented here) of rounds of these three parts results in convergence to a final phasing solution. Within each round, we perform 10 EM iterations, with an additional final EM algorithm run until convergence. This scheme provides the most accurate results and highest log-likelihood.

1. Initialisation of all parameters: ***μ***, *ρ, θ*, **Π**.
2. Repeat until convergence:

(a) thin.
(b) re-phase.
(c) EM iterations.
3. Final EM until convergence.
4. Coancestry curve fitting to estimate dates.

#### Thinning

The thinning refers to a local (gridpoint specific) reduction of the set of possible donors available to copy for each target individual and is a computationally convenient approximation. Briefly, we fit a single layer ancestry unaware model (similar to chromopainter (Lawson *et al*. 2012)) to the full set of donors. For each target individual, we then rank the donors at each gridpoint and pass only the top 100 to the ancestry aware two layer HMM part of our model. For large reference datasets, this greatly reduces the state-space of the model with a negligible reduction in accuracy as typically only a handful of donor haplotypes are likely to be copied from. The reason we do this per individual rather than per target haplotype is to make the donors relevant to both haplotypes available for the re-phasing part of the algorithm. See Appendix Thinning of Donors for full details of how this is done and the impact on computational complexity and accuracy.

#### Re-Phasing

It is important to consider phasing errors as these can potentially mask real ancestry switches or cause our model to infer ancestry switches where there are none. HapMix solves this issue by integrating over all possible phasing at each locus (diploid mode) but this is computationally intensive and Hap-Mix cannot run this option in conjunction with EM inference of the model parameters. To consider all possible phasing is intractable as we would need to consider 2^*G*^ possible phasings per individual genome, where *G* is the number of gridpoints. Instead, we search for phase flips that would lead to an increase in the likelihood of the data under our model and hill climb to a maximum likelihood solution. Although our software can also sample via MCMC other phase solutions, we find that our hill climbing method is both fast and leads to a log-likelihood that is not well improved upon by a long run of the MCMC chain. We refer to our method as “phase hunting”. We use a pass of our fast forward-backward algorithm to see the *marginal* change in log-likelihood under the model were we to flip each gridpoint independently. Examination of these log-likelihood changes informs our phase-hunter as follows:

- Look for spikes in this expected log-likelihood change.
- Flip all of the highest non-overlapping (farther than 0.1cM apart) spikes and refit our HMM.
- Repeat this process as long as the log-likelihood increases.

In practice, we find fewer attractive phase-flips in each successive pass; see Figure 1b.

#### EM Updates

The EM algorithm (sometimes referred to as the Baum-Welch algorithm when used in conjunction with a HMM) performs estimation of the hidden states given a set of parameter estimates (The E-step) and then performs maximum likelihood estimation for the model parameters given these estimates (the M step). Iteration then proceeds over the E and M steps until convergence to a steady set of parameters and local ancestry estimates. At each step the log-likelihood for the model is guaranteed to increase (although convergence to the global maximum is not guaranteed).

### Expected Coefficient of Determination

For simulated data we can report a measure of the accuracy of MOSAIC’s local ancestry. We use the commonly used measure of the squared correlation *r*^2^ between the estimated local ancestry *X* (which is given as the probability that each haplotypes belongs to each ancestry along the genome) and the true local ancestry Z. However, in the absence of such a ground truth *Z* for real data we use an expectation of the accuracy, as per Price *et al*. (2009).

For a given ancestry *a* we compute the expected value of this without knowledge of the true local ancestry *Z_a_* for each individual as follows:

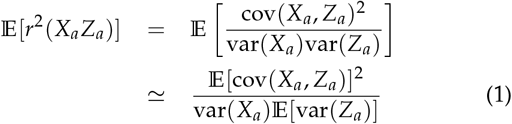

To estimate the numerator in Equation 1 we could take samples from *X_a_* and use the mean of the covariance across the genome between the sampled ancestries and the probabilities *X_a_*. Here we make an approximation as follows:

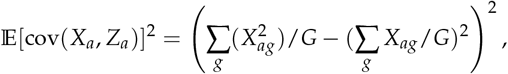

where *X_ag_* is the probability from the model that the individual has ancestry *a* at gridpoint *g* and *G* is the total number of gridpoints. To estimate the denominator in Equation 1 we use

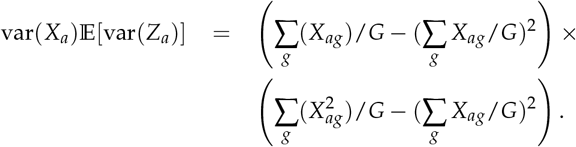

Finally, cancelling some terms we get

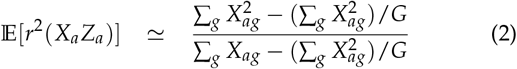

This is then averaged over all ancestries *a* to return the expected squared correlation between the inferred local ancestry and the unobserved true local ancestry.

### F_st_ Summaries

We use F_st_ to summarise the strength of signal and further investigate each admixture event. We first assign *locally* segments of the target chromosomes to the ancestry they are maximally assigned a-posteriori based on the HMM fit. ^2^ These partial, haploid genomes may now be thought of as drifted versions of the “un-admixed” ancestral groups that mixed to create the admixed targets. We then calculate F_st_ (see below) between these so created “un-admixed” genomes and each panel (donor population) used in the model fit.

### F_st_ Calculations

Noting that naïve estimators of 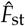 of F_st_ may be biased, and that there are many estimators in the literature, we refer to Bhatia *et al*. (2013) to inform our choice. We follow the recommendations of that paper and use a ratio of averages (rather than an average of ratios). To aggregate across loci, we first define SNP *s* specific 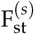 as the estimated variance in the frequency between populations (weighted by population size), divided by the estimated variance of frequency across populations. We can then calculate the genome wide 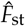 either as the average of such site specific ratios or we can sum the numerator and denominator across the genome first and then find the ratio. Only the latter will converge to the correct underlying value of F_st_ (provided the numerator and denominator are unbiased) and is thus to be preferred.

Note that we deviate from the recommended choice of estimator for the numerator and denominator in Bhatia *et al*. (2013). They recommend the Husdon estimator (Bhatia *et al*. 2013, Equation 5) as this provides unbiased estimates for both the numerator and denominator and is independent of sample size. In fact it assumes equal and large sample sizes. As our sample size for the ancestral groups (constructed as above using maximal assignment ^3^ of local segments of admixed genomes) varies along the genome, we prefer an estimator that is robust to sample size variation and can aggregate site specific contributions to the numerator and denominator with varying sample sizes. We therefore use the popular Weir and Cockerham estimator (Weir and Cockerham 1984) (over 8,000 citations according to Bhatia *et al*. (2013)) in a form that uses a ratio of averages. Specifically we use a variation of Equation 6 in Bhatia *et al*. (2013):

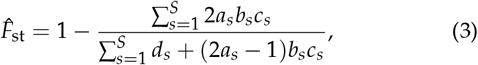

where *s* is the site index, and

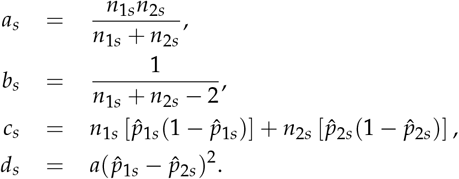

Note that *a* and *b* are unrelated to the *a* and *b* we have used to index ancestries; we are here using the same notation as Bhatia *et al*. (2013) for clarity. Here 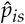 is the observed allele frequency at site *s* for population *i*. The sample size for population *i* is *n_is_* and is fixed for the donor panels but varies for the partial ancestral group genomes created from the admixed haplotypes.

We also calculate 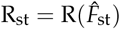 as summarising the estimated average-over-panels relative squared difference in F_st_ between ancestral groups *a* and *b*:

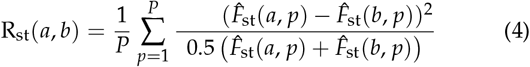

This final statistic is correlated with the estimated 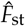 between the latent ancestries (Pearson correlation of 0.23 with p-value of 0.00047 across all real data populations we analysed), however it also takes into account the distances to the donor groups and is thus the more versatile statistic. For example, the mixing groups could be far apart as reported by 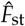 however the panels may be poor surrogates due to drift since admixture. This causes each panel to have a similar 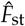 to both ancestral groups then this will show in a low R_st_ overall. The correlation between R_st_ and 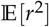 was 0.3 with a p-value of 3.2 × 10^−6^ whereas the the correlation between 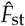 and 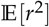 was 0.057 with a p-value of 0.4, showing that Rst is the better predictor of accurate local ancestry.

Finally, it should be noted that when there is no clear admixture signal, MOSAIC returns very small estimated minor ancestry proportions and in this case the estimated F_st_ between the latent ancestries is meaningless. All segments are assigned to one ancestry and the Weir and Cockerham estimator breaks down. R_st_ is still returned but takes values greater than 1. This occurred for four model runs of the extended HGDP dataset (see Extended Human Genome Diversity Panel), all for 2-way admixture; Germany-Austria (GerAus), Karitiana, Lahu, and Welsh. It is noteworthy that GLOBETROTTER also found no strong evidence of admixture for these populations.

### Dating Admixture Events Using Coancestry Curves

We wish to infer the times at which each pair of ancestral groups admixed, based on our model fit. The transition rates matrix Π does not provide a direct estimate of these times (in number of generations), and so we rely on construction of coancestry curves as per Hellenthal *et al*. (2014) with best-fit exponential decay curves to estimate dates. In order to create the coancestry curves in (black lines in Figure 1c) and to find the best fit exponential curve (green lines), we first estimate the probability of being in ancestry *a* at one position and ancestry *b* at a position *d* away, *relative to genome-wide average* probability. This naturally depends on how many generations have occurred since the admixture event. In fact, for two positions *x*_1_ and *x*_2_ *d* apart, this relative probability *P*(*a, b, d*) is approximately equal to *τ_ab_e^−dλ^* + *δ_ab_*. We therefore find constants *τ_ab_, δ_ab_, λ* that minimise the squared difference between *P*(*a, b, d*) and this exponential curve, averaging over all pairs of points these distances *d* apart. We also scale *d* by grid width so that *λ* is in units of generations. We find that the numerical optimisation is relatively unstable so we need to initialise with sensible *τ_ab_, δ_ab_, λ* values. We note that *δ_ab_* ≃ 1 and that at *d* = 0. We also note that *τ_ab_* = *P*(*a, b*, 0) − *δ_ab_* and at any non-zero distance *d* (e.g. where the height of the curve is halfway between the height at 0 and its asymptote)

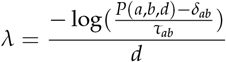

Thus we have crude estimates of *τ_ab_, δ_ab_*, and *λ* with which to start a numerical optimisation routine, which derives the best fit (green lines).

### Data and Code Availability

An open source `R` package is available for download at **https://maths.ucd.ie/~mst/MOSAIC/**. A browser for all results on the extended HGDP dataset (from Hellenthal *et al*. (2014); see Extended Human Genome Diversity Panel) is also provided. The data is a version of publicly available **http://www.hagsc.org/hgdp/files.html** with several additional populations.

## Results and Discussion

### Simulation Studies

Figure 1b and Figure 1c depict a realistic simulation study. We simulated admixture 50 generations ago using real haplotypes from French and Yoruban chromosomes in a 50-50 split; thus the ground truth local ancestry is known. Panels are then formed from Norwegian, English, Ireland, Moroccan, Tunisian, Hadza, Bantu-Kenya (BantuK), Bantu-South-Africa (BantuSA), and Ethiopian individuals. MOSAIC inferred the stochastic relationships between these groups and the underlying mixing groups, along with all other parameters using the automatic algorithm detailed in Algorithm. As can be seen from Figure 1c MOSAIC is able to accurately infer the correct simulated admixture date. Under this simple simulation, the algorithm was also able to infer that one side heavily copies from the West European populations and the other from the African populations.

The current state-of-the-art in local ancestry estimation in the context of a 2-way admixture is provided by HapMix (Price *et al*. 2009). We therefore also ran HapMix in haploid mode with EM to learn its model parameters and then performed a diploid (integration over all possible phasings) run to estimate diploid local ancestry. Reference panels were necessarily provided to HapMix in two sets of proxy haplotypes for the two mixing groups (European and African). This took 50 minutes 5 seconds in comparison with the total MOSAIC run time of 37 minutes 18 seconds (on a standard laptop, which included the time taken to sample the simulated data, fit coancestry curves, etc). Figure 2a shows the true and estimated local ancestries under both models along chromosome 1 for a single diploid individual.

**Figure 2.**
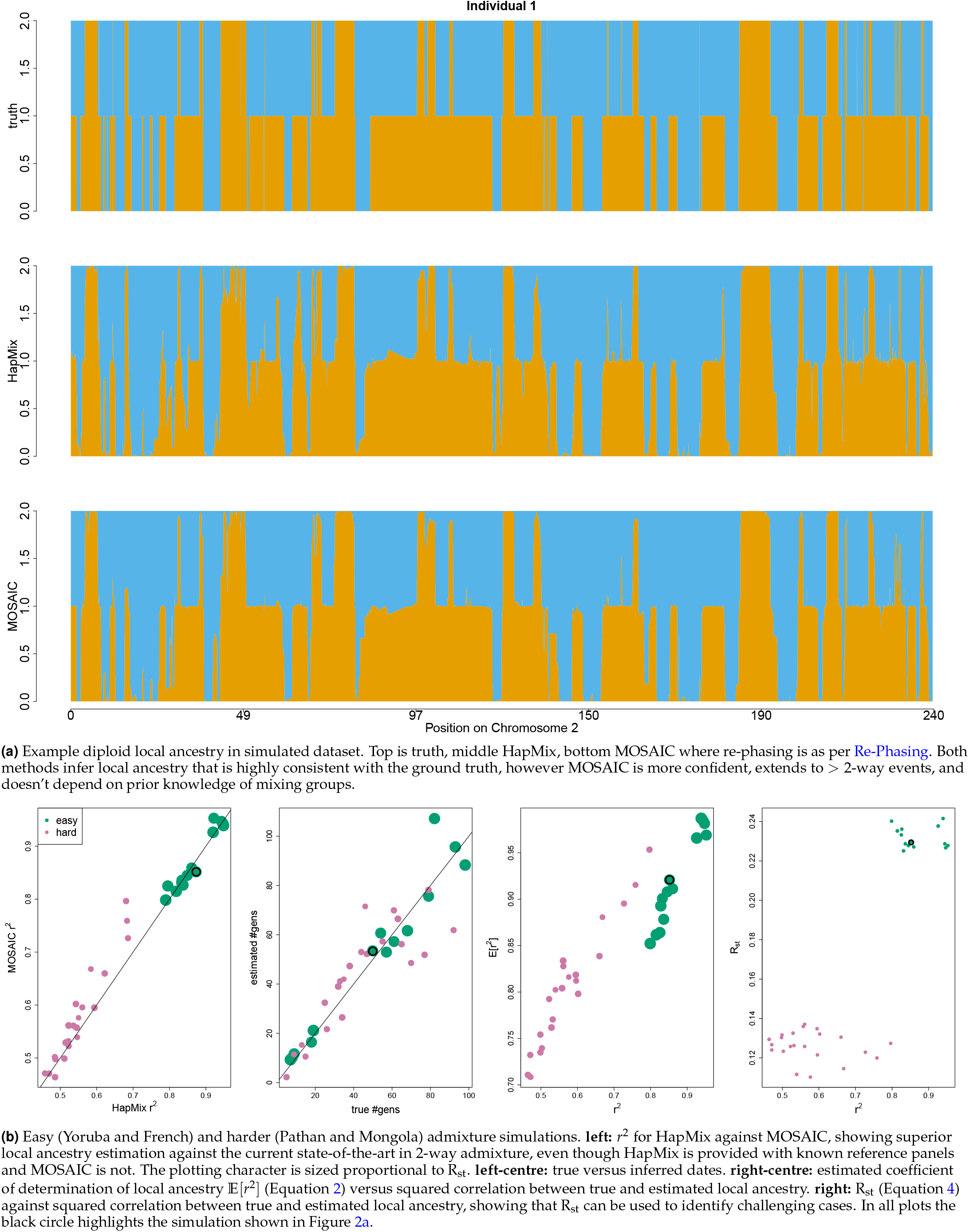
Comparison between MOSAIC and HapMix in 2-way admixture simulations, as described in Simulation Studies.

We next report performance of MOSAIC on scenarios with various levels of difficulty and compare again with HapMix. We fit both models to simulations similar to the above but for varying admixture dates and for a second mixture; the latter is a mixture of Pathan and Mongolan genomes and the reference panels used are Iranian, Lezgin, Armenian, Sindhi, Brahui, Georgian, Hezhen, HanNchina, Han, Tu, Oroqen, Daur, Xibo, Tujia, and Yakut. Again, these must be provided to HapMix as two sets of known surrogates whereas MOSAIC infers the relationships to the admixed genomes. See Figure 2b for a summary of MOSAIC’s and HapMix’s performances across these scenarios. MOSAIC typically outperforms HapMix in terms of *r*^2^ with the true local ancestry, especially for the more difficult scenarios.

It must be stressed here that this is a setting that is ideal for HapMix (2-way admixture with known, highly appropriate reference panels) whereas MOSAIC generalises to multi-way admixture and can even thrive when the reference panels are not good surrogates for the mixing ancestral groups. Furthermore, MOSAIC infers the stochastic relationships between panels and ancestries.

### Extended Human Genome Diversity Panel

#### Two-way, Single Event

MOSAIC handles multiway admixture and provides accurate local ancestry inference, however we first restrict to the case of 2-way admixture events in order to compare results with the current state-of-the-art method GLOBETROTTER (Hellenthal *et al*. 2014). Figure 3 shows the inferred dates for the target populations that were found to have experienced a single 2-way admixture event according to Hellenthal *et al*. (2014). The x-axis shows the inferred dates (as generations since admixture) along the with 2 standard errors and the y-axis shows the MOSAIC inferred dates.

**Figure 3.**
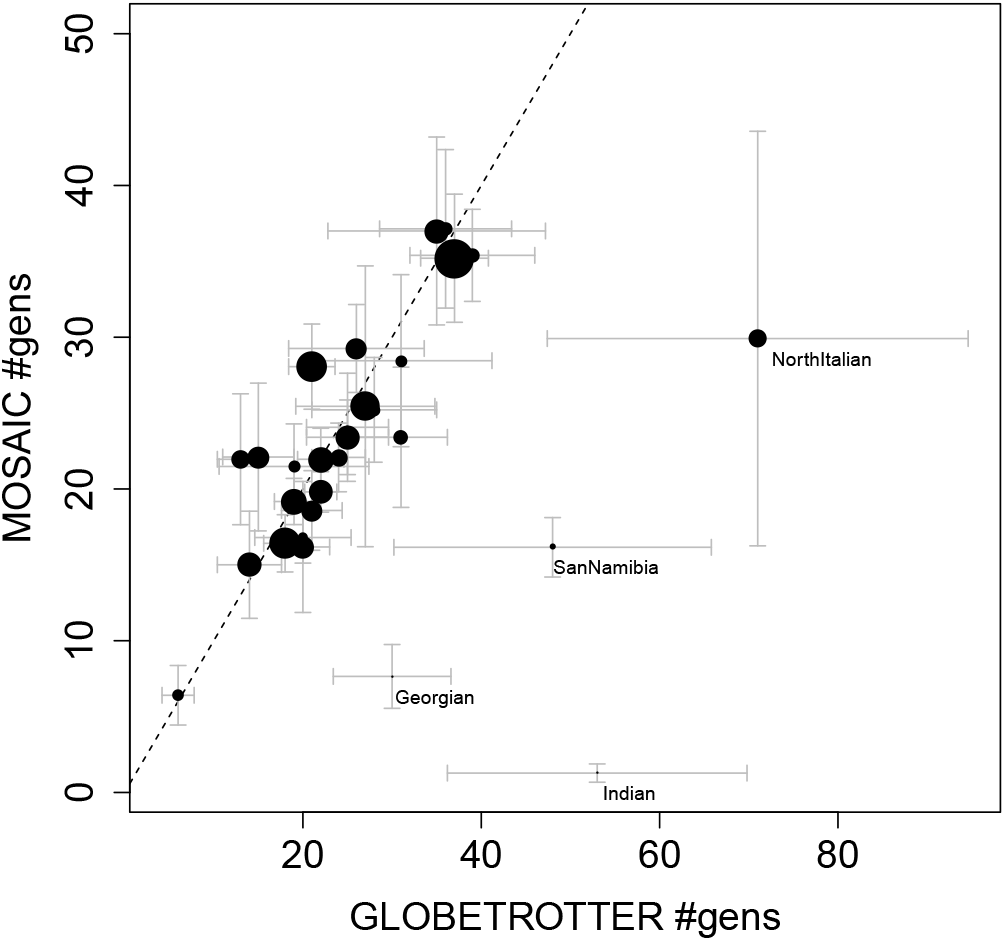
Inferred dates from MOSAIC are plotted against inferred dates from GLOBETROTTER, including bootstrapped ±2 standard error bars for real data 2-way admixture events (see note in Two-way, Single Event). The GLOBETROTTER dates are from Table S14 of Hellenthal *et al*. (2014). The size of the central disc of each event is proportional to R_st_ (see F_st_ Summaries). Note that Melanesian is not shown as MOSAIC infers a very old mixing event of 221 generations ago.

The GLOBETROTTER paper creates bootstrapped *chromosomes* and finds the sample distribution of dates based on these, a protocol we follow here. Note that although MOSAIC models a single admixture event that may be experienced at different times by different admixed individuals we here fit a set of coancestry curves with a common rate parameter for consistency with Hellenthal *et al*. (2014) and to provide a direct comparison, In Figure 3 most points fall close to the 1 – 1 line, showing a strong agreement between the two approaches. MOSAIC provides tighter bootstrapped confidence intervals on average but we observe that there are some targets for which MOSAIC infers a far more recent admixture with very narrow confidence intervals (bottom right of Figure 3). We note that several warning flags are raised when we analyse the output from MOSAIC for these three populations for a 2-way admixture event.

The R_st_ statistic (which is based only on the MOSAIC results) is smallest for the target populations that have the strongest disagreement with GLOBETROTTER (Georgia, San-Namibia, and India; labels appear below and to the right of plotting points in Figure 3 for these populations). For India, Sindhi is the closest (smallest 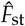 to **both** admixing groups. Similarly, for San-Namibia the San-Khomani are the best match to both ancestries. In the Georgian case, the Armenians and Russians are extremely close in 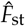 to both ancestries. Note that for the Georgian target in Hellenthal *et al*. (2014), ROLLOFF (Patterson *et al*. 2012) failed to run without error. This method is more suited to very old admixture events and does not require the ability to infer admixture breakpoints along the genome. In the GLOBETROTTER analysis of India, the major ancestry is represented by South Central Asian populations (Sindhi, Pathan, Indian Jew), however the minor ancestry (14%) is highly diverse, consisting of Asian populations (Cambodian, Mongola, Han) as well as Ethiopian and Papuan. We find a similar 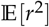 for 2 and 3 way admixture (0.604 and 0.601 respectively), although neither exhibit a clear admixture signal as measured by the R_st_ statistic (0.0045 and 0.016 respectively).

Interestingly, in Supplementary Section 6.4.5 of Hellenthal *et al*. (2014), the authors compare two closely related methods for dating admixture; using coancestry curves (as per Dating Admixture Events Using Coancestry Curves but in the absence of an explicit model for the latent ancestry, applied to each pair of the donor groups) and a second method wherein these coancestry curves are standardized by those of a “NULL” individual. To create this individual, “across-individual” coancestry curves are generated by taking painting samples from *different* individuals. These are then used to normalise the original coancestry curves. Simulation studies in that paper show that this has the effect of eliminating spurious signals of admixture, leading to more reliable date estimates. However inference of source groups is negatively impacted. The authors note that “we expect both approaches to generate coancestry curves that largely agree in their conclusions, in particular their date estimates, and disagreement might signal lower reliability of the results”. For the 27 target populations analysed in this section, they find that there are two populations wherein the two dating procedures “differ enough that it may substantially affect interpretation of the events … the Melanesian and the Namibian San”. Indeed the GLOBETROTTER results for Melanesia are confusing; the “Papuan-like” ancestry is represented by some Asian groups - Papuan (21.0%), Cambodian (3.0%), Sindhi (2.6%), and Japanese (2.5%) - but also such diverse groups as Turkish (1.5%), Uygur (1.3%), Indian (1.3%), Moroccan (1%), Greek (1%), and Uzbekistani (Uzbek) (1%). Conversely, MOSAIC obtains a stronger signal of 3-way admixture according to 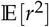 (0.75 versus 0.63 for 2-way). Even then the 2-way result is less ambiguous, with a clear African minor ancestry and South-East Asian plus Papuan major ancestry. The 3-way result splits the Papuans off from the rest of South-East Asia. 2-way MOSAIC finds a very old signal of admixture for the Melanesians of 221 generations, however there does also appear to be a very recent admixture event also.

We now turn our attention to a number of case-studies of high confidence, specifically Hazara, Bedouin, and Chuvash 2-way, Maya 3-way, and San-Khomani 4-way admixture models. Figure 4 depicts the inferred copying matrix ***μ*** for each of these case-studies and Figure 5 shows the coancestry curves used to date the events. Note that the bootstrapped standard errors for the dates in Figure 3 are created using a single rate parameter estimate whereas the plots shown here estimate one such parameter for each unique unordered pair of ancestry (i.e. 1 – 1,1 – 2,2 – 2 for 2-way admixture). For a 3-way (or higher) model, a single date is invalid and we therefore provide one such coancestry plot for each pair of ancestries in Figure 5.

**Figure 4.**
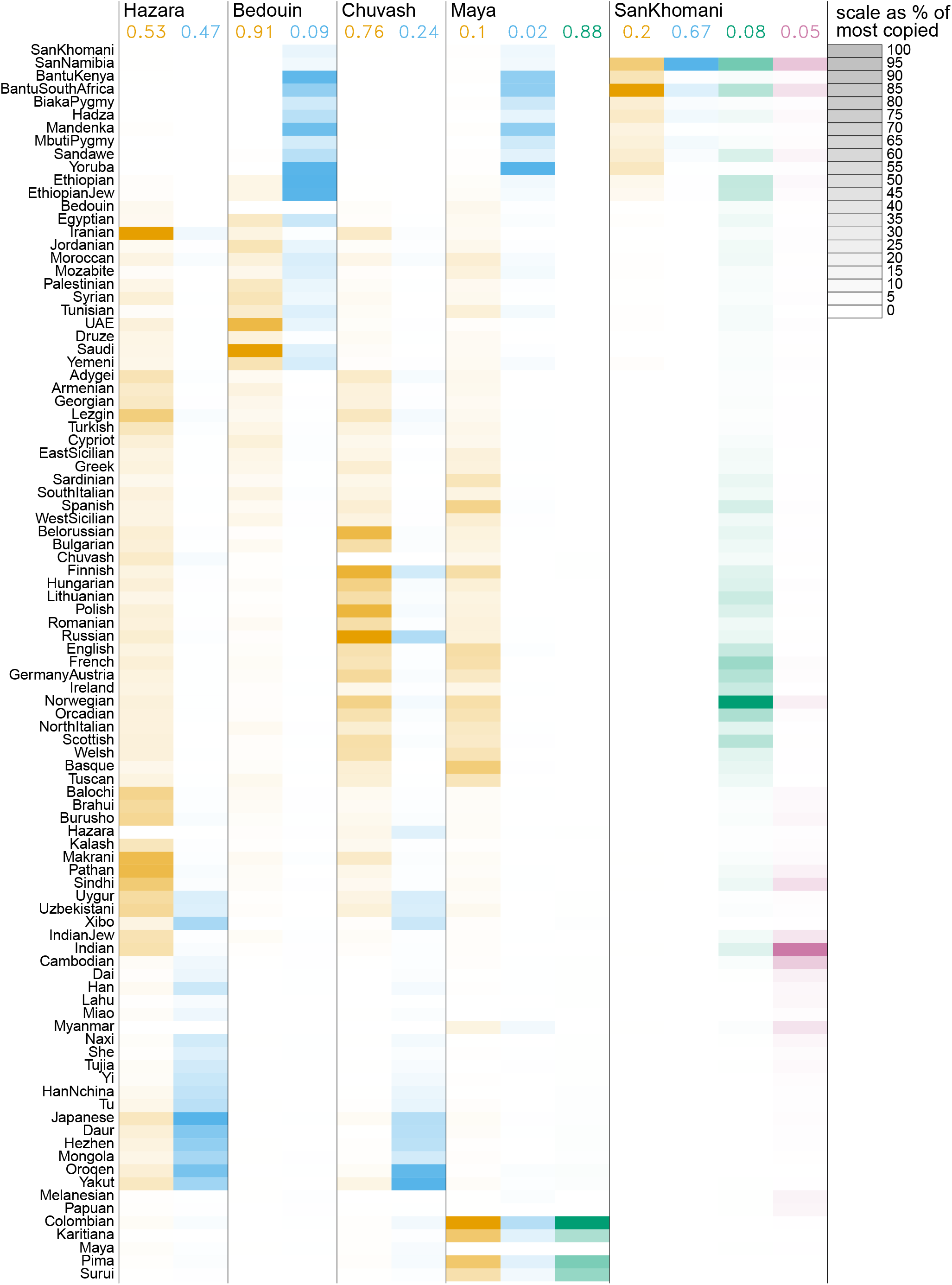
Inferred copying matrices for case studies of human admixture based on the HGDP dataset. The copying proportions ***μ***_*pa*_ are scaled within columns to % of the most copied donor population so that each cell shading is equal to 100.***μ***_*pa*_ arg max_*p*_ ***μ***_*pa*_. Along the top are the marginal ancestry proportions for each admixed target.

#### Hazara 2-way

Figure 6 illustrates the accurate local ancestry estimation available for a recent admixture between 2 well diverged groups. Hellenthal *et al*. (2014) find that the Hazara “show the clearest signal of admixture in the entire dataset”; this is reflected by MOSAIC inferring tightly coupled coancestry curves (Figure 5a) with highly confident local ancestry estimates (Figure 6) and well diverged mixing groups (Table 1). We have chosen to include local ancestry estimates for two individuals along two chromosomes, but these are representative of the entire population.

**Figure 5.**
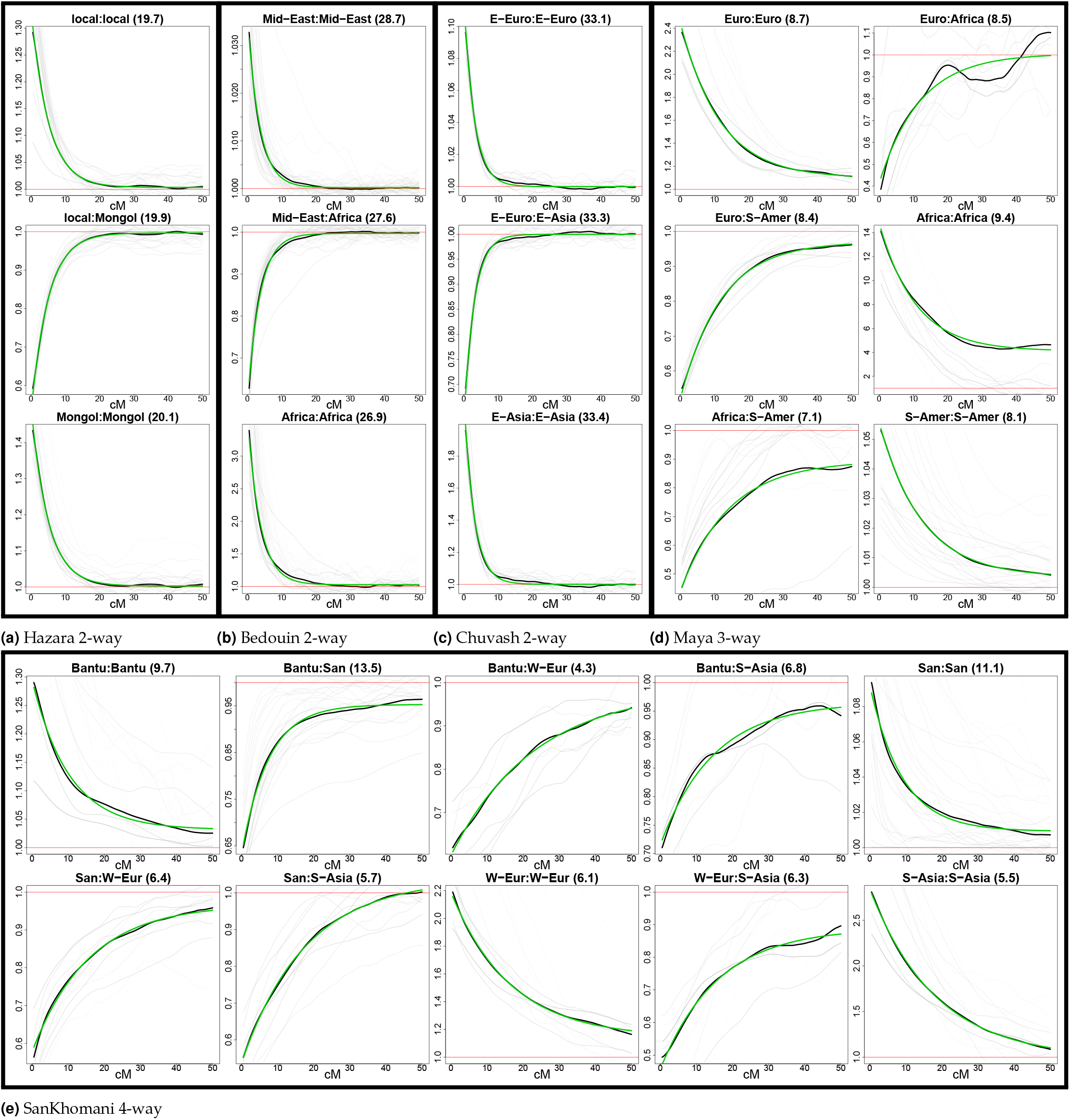
Coancestry curves for case studies of admixture within the HGDP dataset. On the top of each sub-plot the ancestry sides are labelled according to the closest donor panel as measured by 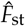 (see Tables 1 to 3) and the estimated number of generations since admixture between each pair of ancestries is given in brackets.

**Figure 6.**
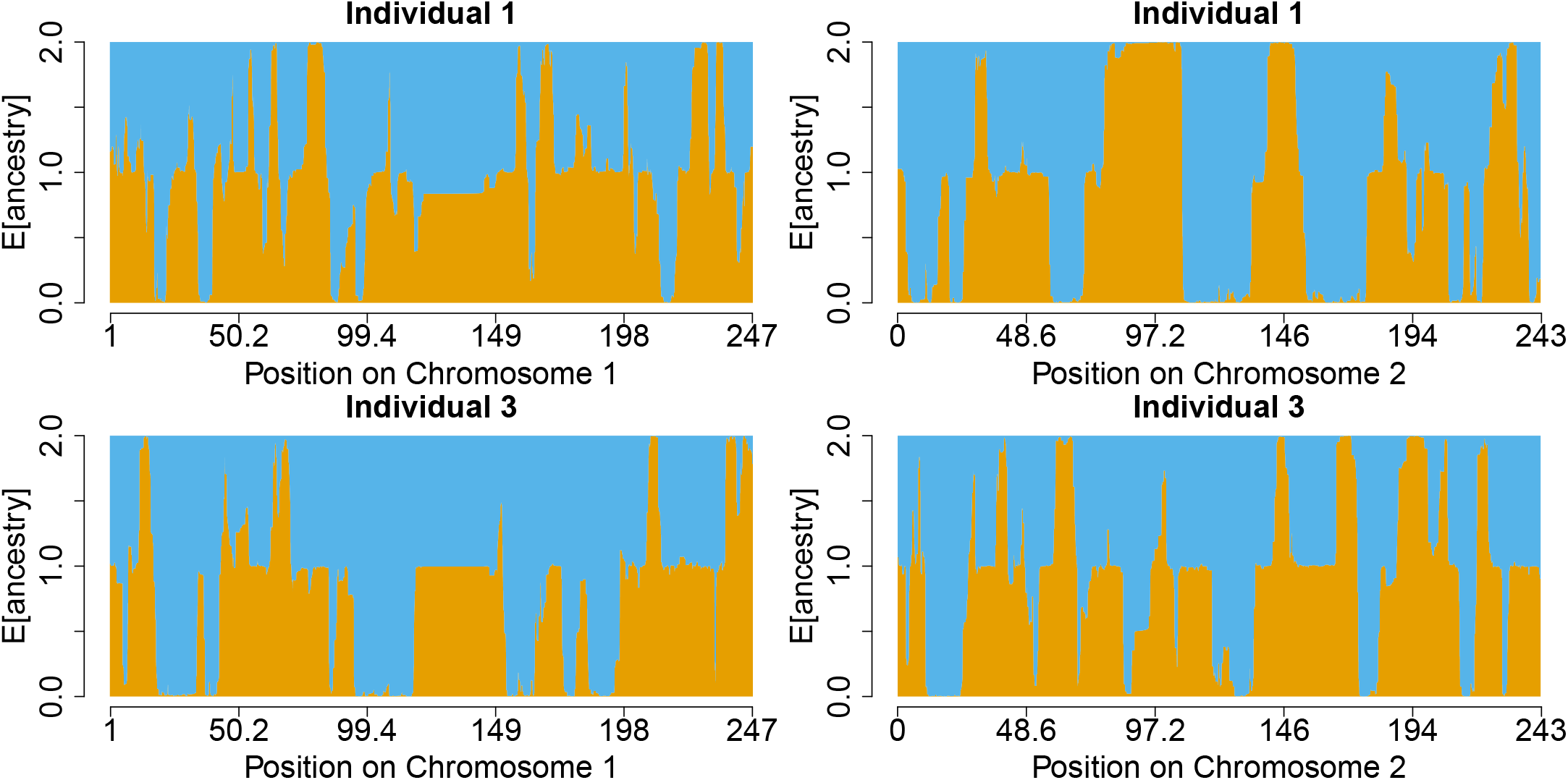
Hazara estimated local ancestry on chromosomes 1 and 2 across two individuals. There is a roughly 50-50 ancestry contribution from two sources approximately 20 generations ago. The orange source is Pathan like and the blue is Mongolian type (see Table 1 for details). The colours are consistent with Figure 4 which shows scaled copying proportions for each donor panel.

**Table 1.** F_st_ estimates between local ancestries and the closest 5 panels in Hazara 2-way. 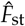 between the inferred local ancestries is 0.087. R_st_ is 0.093, showing substantial divergence between the mixing groups and good representation of them using modern populations.

|  |  |  |  |
| --- | --- | --- | --- |
| Pathan | 0.0066 | Mongola | 0.0075 |
| Iranian | 0.007 | Xibo | 0.0084 |
| Turkish | 0.0078 | Daur | 0.0089 |
| Balochi | 0.0089 | Oroqen | 0.01 |
| Sindhi | 0.0089 | Hezhen | 0.011 |

#### Bedouin 2-way

As we present results of applying MOSAIC to multiple North African populations with some shared history in Selection Signal at the HLA in North Africa, we first include MOSAIC results on one of these populations. The Bedouin are a nomadic group that are well characterised by an admixture between sub-Saharan and Middle Eastern groups.

**Table 2.**
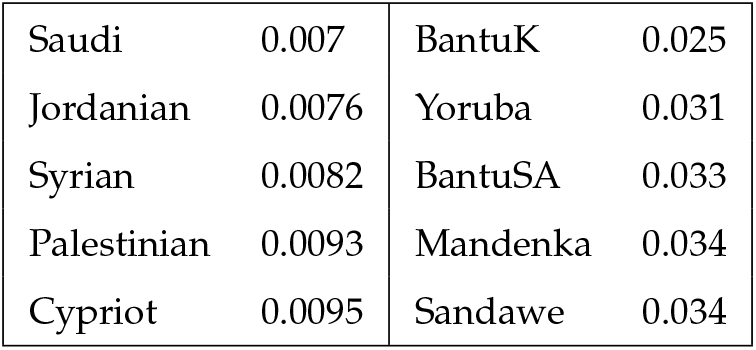
F_st_ estimates between local ancestries and the closest 5 panels in Bedouin 2-way. 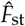 between the inferred local ancestries is 0.13. R_st_ is 0.11, showing substantial divergence between the mixing groups and good representation of them using modern populations.

|  |  |  |  |
| --- | --- | --- | --- |
| Saudi | 0.007 | BantuK | 0.025 |
| Jordanian | 0.0076 | Yoruba | 0.031 |
| Syrian | 0.0082 | BantuSA | 0.033 |
| Palestinian | 0.0093 | Mandenka | 0.034 |
| Cypriot | 0.0095 | Sandawe | 0.034 |

#### San-Khomani 4-way

As per Choudhury *et al*. (2017) we find evidence of admixture between Bantu-speakers, Khoesan, Europeans, and populations from the southern Asia. In fact the Choudhury *et al*. (2017) authors conclude that “the absence of population-scale WGS data for KS groups restricted our ability to fully utilize our WGS data in analyses such as admixture mapping and local ancestry detection. The availability of such data would enable a more comprehensive analysis and is expected to provide novel insights.”. A strength of MOSAIC is that it finds haplotypes from single sources embedded within admixed donor genomes and can then disambiguate the mixing sources. There is confounding between the only donor panel with a significant Khoesan component (San-Namibia, who have a similar admixture history to the San-Khomani) and the Bantu-type donors, which manifests in the copying matrix ***μ*** exhibiting a large preference for copying San-Namibia haplotypes within **both** of these African-type ancestries; however the F_st_ based analysis is able to separate the actual haplotypes involved so that a clear disambiguation between all 4 ancestry components is achieved (see Table 3). See also the Supplementary document for a 3-way admixture analysis that combines the two African ancestry types.

**Table 3.** F_st_ estimates between local ancestries and the closest 5 panels in SanKhomani 4-way. The F_st_ estimate between the inferred local ancestries is 1×2=0.1 1×3=0.14 1×4=0.14 2×3=0.27 2×4=0.27 3×4=0.066. The R_st_ is 1×2=0.057 1×3=0.12 1×4=0.078 2×3=0.24 2×4=0.2 3×4=0.026.

|  |  |  |  |  |  |  |  |
| --- | --- | --- | --- | --- | --- | --- | --- |
| BantuSouthAfrica | 0.0066 | SanNamibia | 0.02 | French | 0.011 | Uygur | 0.024 |
| BantuKenya | 0.01 | BiakaPygmy | 0.09 | Welsh | 0.011 | Uzbekistani | 0.025 |
| Yoruba | 0.012 | BantuSouthAfrica | 0.09 | Bulgarian | 0.012 | Sindhi | 0.025 |
| Mandenka | 0.018 | Sandawe | 0.099 | Romanian | 0.012 | Hazara | 0.026 |
| Sandawe | 0.022 | MbutiPygmy | 0.11 | Spanish | 0.012 | Indian | 0.027 |

Figure 7a illustrates the accurate local ancestry estimation available for such a recent admixture between 4 well diverged groups. We have chosen to include local ancestry estimates for three individuals with markedly different ancestry components:

2 2-way admixture between Bantu 32% and Khoesan 68%.
1 3-way admixed Bantu 45%, Khoesan 51%, and European 4%.
19 4-way admixed Bantu 13%, Khoesan 45%, European 26%, and Asian 16%.

**Figure 7.**
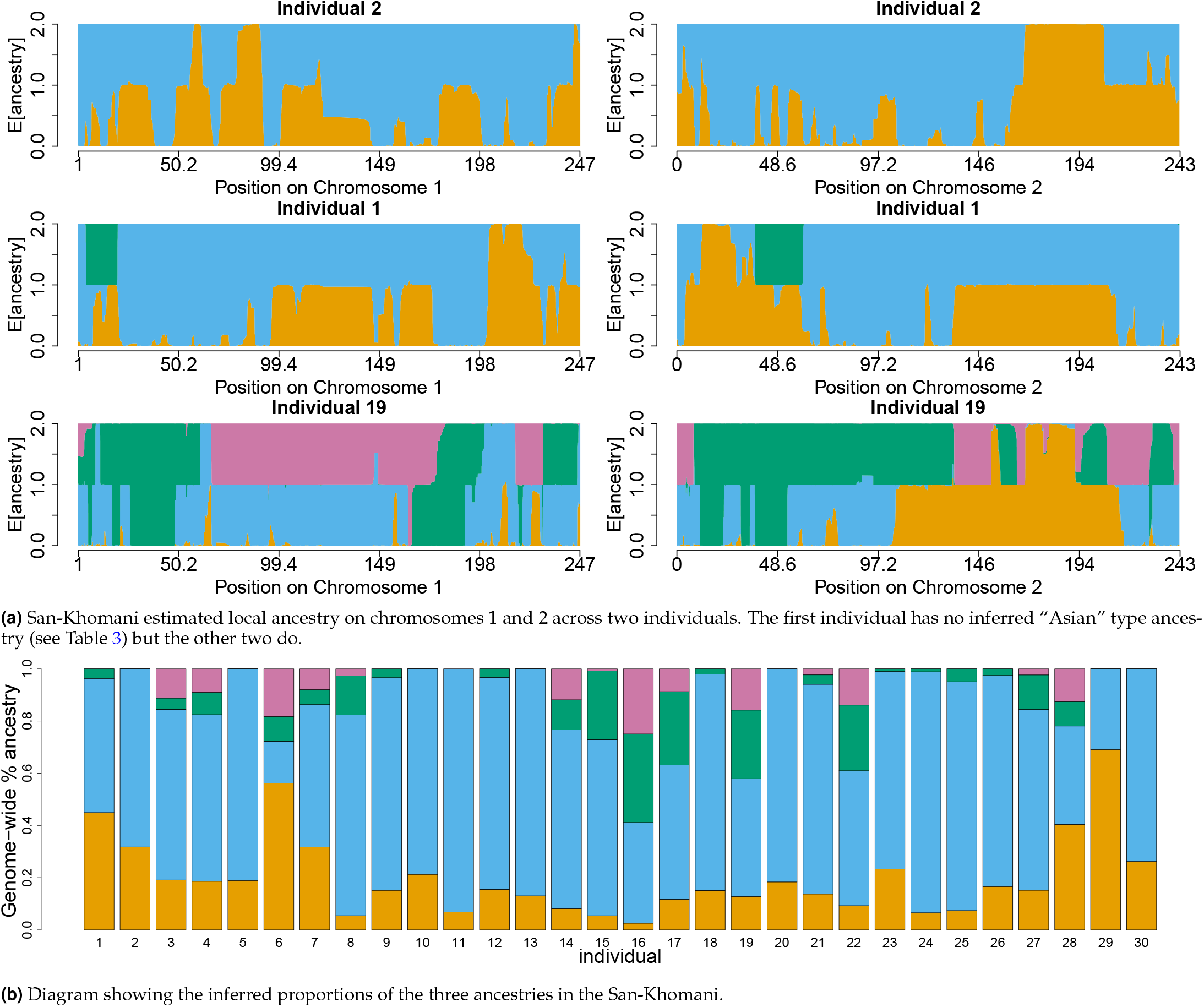
Details of San-Khomani 4-way admixture model fit. The orange source is Bantu-like, blue is San, green is European, and purple is Asian (see Table 3 for details). The colours in both plots are consistent with Figure 4 which shows scaled copying proportions for each donor panel.

This illustrates the flexibility of the method as it successfully infers the variable mixing proportions across individuals in a single model fit. Note that although Indian is the most copied source for the Asian side (scaled by number of donors, see Figure 4), the closest donor panel in 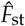 to the mixing source is in fact Uygur (see 4th column of Table 3); however Uzbekistani, Sindhi, Hazara, Indian, Pathan and Burusho all score similarly close in terms of genetic divergence to this unseen South Asian ancestral group.

### Selection Signal at the HLA in North Africa

We demonstrate a test for selection of alleles as manifested by an excess in one ancestry over another in a genomic region, when averaged over multiple individuals experiencing the same admixture history. We see a large spike in African versus European ancestry at the Human Leukocyte Antigen (HLA as per Zhou *et al*. (2016), and a excess of IBD sharing as per Botigué *et al*. (2013)) when we allow MOSAIC to use a small set of reference panels from Europe and sub-Saharan Africa. When we allow MOSAIC to infer which panels to use and how, the spike is greatly reduced, due to preferential copying of African type HLA alleles extant in Southern European populations.

The HLA spike represents the highest and widest segment of significantly different average ancestry (as measured by a simple z-test of local average African ancestry proportion, compared to the genome-wide average and standard error based on genome-wide standard deviation; Figure 8a lower and Figure 8b lower). The spikes in mean African ancestry that are at 1*p*, 6*q*, etc are narrow and therefore unlikely to be a signal of post-admixture selection as the event is dated to 31 generations ago as simulations of positive selection at a single site following admixture between sub-Saharan African genomes and European genomes with the same mixing proportions and the same time since admixture resulted in broader spikes (see Appendix: Simulation of positive selection). Further tests in support of this post-admixture hypothesis are provided in the supplementary document.

**Figure 8.**
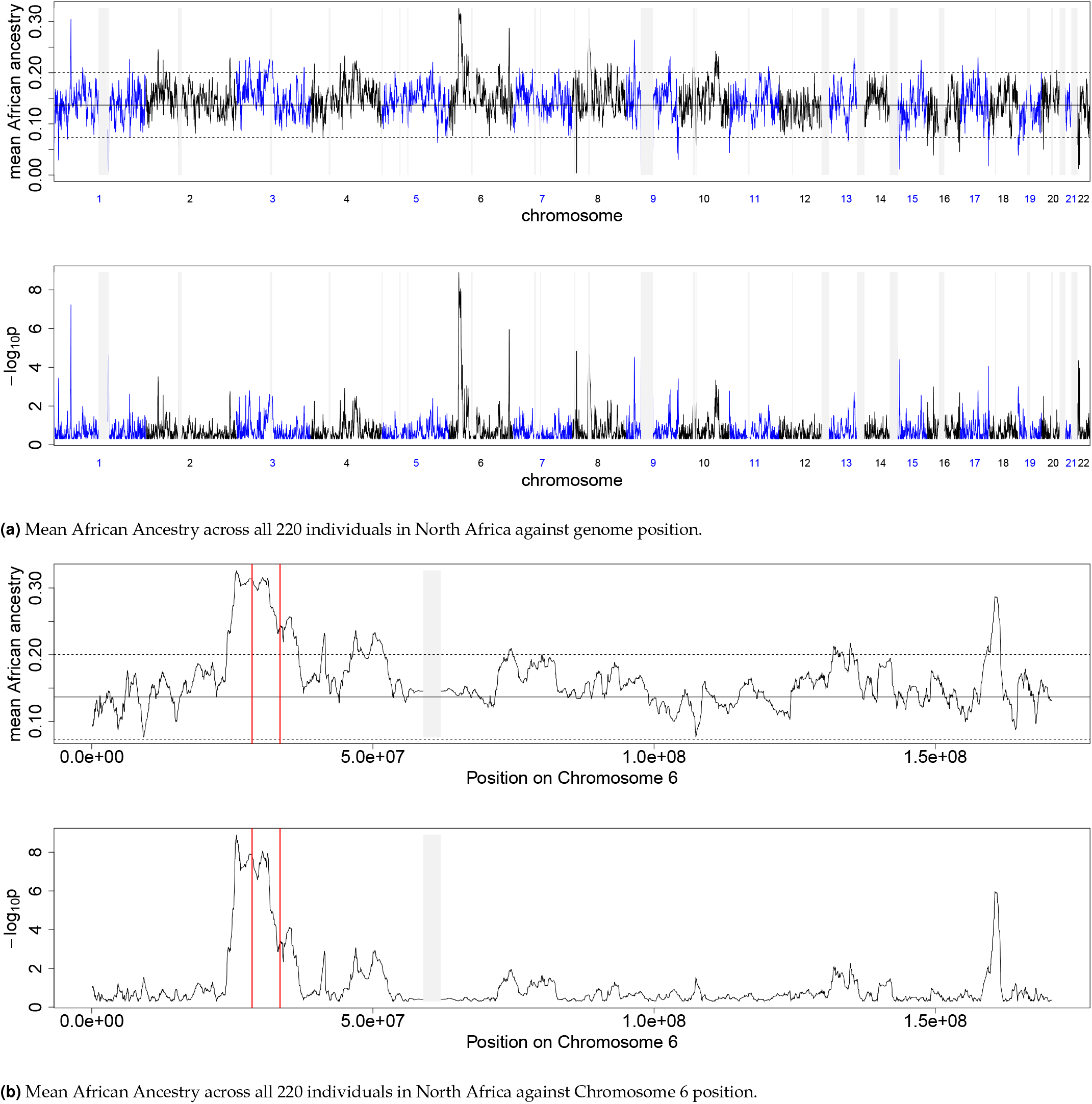
Mean African Ancestry. There is a high and wide spike at the HLA on Chromosome 6 at the HLA. Note that we have blocked out (in light grey) all 1Mb regions with fewer than 10 markers; this includes centromeres with low recombination rates and few SNPs.

## Discussion

We have created freely available software that implements an efficient and highly accurate model for fitting multi-way admixture events. We call this method and the associated software MOSAIC. It can not only handle multi-way admixture events but, unlike all currently available methods, it infers the stochastic relationship between groups of potential donors to use in the Li and Stephens’ type Hidden Markov Model and the underlying ancestral groups.

We have demonstrated that even in the case of two-way admixture we can more accurately estimate both local ancestry along the genome and the parameters governing the event (number of generations since admixture and proportion of mixing ancestries) than the current state-of-the-art approaches.

## Acknowledgements

This work was funded by the NIH (grant R01 HG006399) and the Wellcome Trust (project number H5RXUH00). The extended HGDP dataset was cleanly provided by George Busby. Fertile conversations that helped develop ideas for this work were contributed to variously by Daniel Falush, Daniel Lawson, Garrett Hellenthal, and Alkes Price.

## Appendix: Computational Efficiencies

### Unique Haplotype Lists

In order to reduce the memory footprint of the model and algorithm, we store only the unique haplotypes at each gridpoint for the donors and target admixed genomes. These lists grow sub-linearly with the number of samples and size of reference panels so that the method scales to huge datasets. We first read in the data SNP by SNP; each locus is assigned to the nearest gridpoint so that some gridpoints have several, some one, and some no observations. The emission probabilities simply require knowledge of the number of matches between any potential donor and each target and the number of observations of each at that gridpoint. This is stored efficiently in a table for fast lookup as required i.e. we require a table of all possible values of *θX* and (1 – *θ*)(*W – X*) where *θ* is the estimated copying error rate, *X* is the number of mis-matches and *W* is the number of observations. We enumerate all possible values of *X* and *W* once after compression and update the lookup table (which is modest in size, depending on the density of markers and number of individuals in the study) after each re-estimation of *θ* in our EM algorithm. A map from (donor, recipient, gridpoint) to this lookup table is also constructed exactly once to complete the compression to a grid. We obtain lossless compression of the raw genotype data on the recombination grid to between 1 and 2 percent for each target population in the HGDP dataset when taking all other populations as potential donors using this scheme.

### Thinning of Donors

The total number of hidden states in our two layer HMM is *A* × *N* where *A* is the number of admixing populations and *N* is the total number of donor haplotypes. However, in practice, the forward-backward algorithm will place a non-negligible probability on only a small subset of the donors at any one position. We exploit this fact for computational efficiency as follows:

1. Fit the ancestry unaware single layer HMM using all potential donor haplotypes.
2. Use this to *rank* the donors *at each gridpoint* from highest to lowest forward-backward probability.
3. Record the vector of top *ng* such donors at each gridpoint for each haplotype such that over 99% of the probability is captured, up to a maximum of 100 donors.
4. Create the superset of the donors for both haplotypes of each individual at each gridpoint.
5. Use only these locally thinned set of donors at each grid-point in the full two-layer ancestry aware HMM.

We use the top ranked donors for both haplotypes for each individual so that the thinning does not overly effect the phasing (see Section Re-Phasing)). i.e. both haplotypes can see and use the potential donors of the other haplotype belonging in each target individual. We find that using fewer than 100 of these local best donors gave results almost identical to allowing local copying from any of the full set of donors. For example, the median number of donors required to capture more than 99% of the copying probability in the ancestry unaware model was 66 for admixture simulated between French and Yoruban genomes and 59 for admixture between French and English.

Thinning in this way means that the ancestry aware parts of the algorithm scale *independently* of the number of donors in the dataset. The thinning steps scale linearly with the number of donors and independently of the number of hidden ancestries, but these steps need only be performed several times in total (Section Algorithm). Once the HMM states have been so reduced, we need to track locally at each gridpoint which donors occur where in the thinned set, one gridpoint to the left (forward algorithm) and one gridpoint to the right (backward algorithm) as we need to calculate the probability of switches to the same haplotype. We therefore include two additional lists of donors at each gridpoint; the indices of the matching donors (should they appear in the top 100) to the left and to the right of each gridpoint.

### Computational Tricks for the HMM

In the forward (and backward) algorithms at each gridpoint, we need to compute the probability of switching from each hidden state into every other hidden state which is 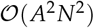. However, as per Fearnhead and Donnelly (2001) and Li and Stephens (2003), we do not need to examine all pairs of donor haplotypes. All switches are equally likely (given the latent ancestry switch status) except the non-switch where the same haplotype is copied for consecutive gridpoints. Thus when examining the possible switches between gridpoint *g* – 1 and *g*, we only need sum over all probabilities for copying any donor haplotype at *g* – 1 and multiply our transition and emission probabilities by this with a small correction for self-switches.

### Initialisation

To initialise our model, we first fit a single layer HMM that is ancestry unaware to the full set of donors genomewide. This is similar to a gridded version of chromopainter (Lawson *et al*. 2012). We perform EM until convergence to estimate *ρ, θ*, **M**, where the latter is simply a vector of copying probabilities for each group.

We now wish to initialise values for the ancestry aware part of the model i.e. an **M** copying matrix with one column per hidden ancestry and the marginal probabilities of each ancestry *α*. We first impose windows of size 0.5*cM* across the genome and calculate the expected number of switches into each donor group in each window. This window size is chosen so that there is typically one latent ancestry per window but multiple donor groups are copied from (the number of generations undergoing recombination until we expect a single event in a window is 200). We then use EM to fit a mixture model where the number of mixtures is the number of hidden ancestries we wish to model. For haplotype *h* in window *w*, the expected number of switches 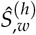 into the panels is modelled as a mixture of Multinomials.

## Appendix: Comparison with GLOBETROTTER for 2-way admixture events

As we regard GLOBETROTTER to be both the closest in spirit to our approach (in terms of genome-wide estimate) and the state-of-the-art in estimating time since admixture we expand upon the comparison provided in Section Two-way Single Event. Table 4 compares results for the two-way, admixture events inferred from the expanded HGDP dataset and comprises the values used to create Figure 3. The contribution of our method lies in accurate multiway local ancestry estimation within a single model and framework that provides these estimates.

**Table 4.** Comparison of date estimates from MOSAIC and from GLOBETROTTER for all inferred 2-way admixed populations in the extended HGDP dataset

| Population | n | source 1 | source 2 | Rst | GLOBETROTTER | MOSAIC |
| --- | --- | --- | --- | --- | --- | --- |
| Hazara | 22 | Pathan | Mongola | 0.0931 | 22 ± 0.9 | 20 ± 0.7 |
| Uzbekistani | 15 | Turkish | Mongola | 0.102 | 19 ± 1.1 | 19 ± 0.8 |
| Uygur | 10 | Iranian | Mongola | 0.101 | 22 ± 1.3 | 22 ± 1 |
| Makrani | 22 | Balochi | BantuKenya | 0.124 | 18 ± 1.2 | 16 ± 0.9 |
| Druze | 42 | Cypriot | Ethiopian | 0.159 | 37 ± 1.9 | 35 ± 2.1 |
| Mozabite | 25 | Moroccan | Yoruba | 0.122 | 21 ± 1.3 | 28 ± 1.4 |
| Turkish | 17 | Armenian | Uygur | 0.0662 | 24 ± 1.5 | 22 ± 1.1 |
| Brahui | 23 | Balochi | Ethiopian | 0.0885 | 20 ± 1.5 | 16 ± 2.2 |
| Yemeni | 4 | Jordanian | Ethiopian | 0.0954 | 14 ± 1.8 | 15 ± 1.8 |
| Pima | 14 | Egyptian | Maya | 0.0422 | 6 ± 0.9 | 6 ± 1 |
| BantuSouthAfrica | 8 | SanKhomani | Yoruba | 0.0113 | 25 ± 2.3 | 24 ± 1.8 |
| Tu | 10 | Turkish | HanNchina | 0.0947 | 25 ± 2.3 | 23 ± 1.2 |
| WestSicilian | 10 | EastSicilian | Ethiopian | 0.117 | 27 ± 3.9 | 25 ± 4.6 |
| Cambodian | 10 | Uygur | Dai | 0.035 | 20 ± 2.7 | 17 ± 0.8 |
| Georgian | 20 | Armenian | Russian | 0.00338 | 30 ± 3.3 | 8 ± 1.1 |
| Romanian | 13 | Bulgarian | Uygur | 0.0545 | 31 ± 2.6 | 23 ± 2.3 |
| Bulgarian | 18 | Romanian | Uzbekistani | 0.0433 | 28 ± 3.5 | 25 ± 1.7 |
| Hezhen | 8 | Uzbekistani | Daur | 0.0711 | 13 ± 1.3 | 22 ± 2.2 |
| Oroqen | 9 | Uzbekistani | Daur | 0.0855 | 15 ± 2 | 22 ± 2.4 |
| Hungarian | 18 | GermanyAustria | Uygur | 0.0535 | 39 ± 3.5 | 35 ± 1.5 |
| HanNchina | 10 | Uzbekistani | Tujia | 0.082 | 26 ± 3.8 | 29 ± 1.4 |
| Daur | 9 | Turkish | Mongola | 0.0814 | 21 ± 1.7 | 19 ± 1.3 |
| Greek | 20 | EastSicilian | Ethiopian | 0.0493 | 36 ± 3.7 | 37 ± 2.6 |
| Melanesian | 10 | Sandawe | Papuan | 0.0475 | 28 ± 7.6 | 221 ± 41.7 |
| Mandenka | 22 | Yoruba | Ethiopian | 0.0438 | 19 ± 4.2 | 21 ± 1.4 |
| Indian | 13 | Sindhi | Sindhi | 0.00332 | 53 ± 8.4 | 1 ± 0.3 |
| NorthItalian | 12 | Spanish | Tunisian | 0.0694 | 71 ± 11.8 | 30 ± 6.8 |
| Polish | 16 | Belorussian | Uzbekistani | 0.0433 | 31 ± 5.1 | 28 ± 2.8 |
| Tuscan | 8 | WestSicilian | Ethiopian | 0.0954 | 35 ± 6.1 | 37 ± 3.1 |
| SanNamibia | 5 | SanKhomani | SanKhomani | 0.0187 | 48 ± 8.9 | 16 ± 1 |

## Appendix: Simulation of positive selection

We simulated positive selection in an admixed population for a single locus on Chromosome 6. We begin with ancestry proportions equal to those inferred in North Africa (minor ancestry of 15%, see Section Selection Signal at the HLA in North Africa). Using *N_e_* = 10, 000 diploid individuals, we simulated random recombinations along chromosomes of genetic length equal to Chromosome 6 (1.93 Morgans) for 31 generations. The number of recombinations is Poisson with rate 1.93 and the locations of the recombinations are uniform along genetic distance. We assume a Wright-Fisher with selection model of random mating amongst individuals and keep track of the simulated ancestry switch points continuously along each chromosome, as well as the ancestry of each segment. We set a non-zero selection coefficient at a single locus of *s* = 0.035 such that haplotypes containing ancestry of the minor type *a* at this locus are up-weighted with a relative weight of 1 + *s*. i.e. when considering parents, each individual selected parents randomly with the probability of selecting a parent of ancestry *a* given by 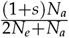, where *N_a_* is the number of haplotypes of ancestry *a* at the selected locus. After 31 generations, we have 10^4^ admixed individuals from which we sub-sample 220 diploid individuals (440 haplotypes). We then plot the average ancestry across these individuals against locus in Figure 9, noting the spike centred at the locus simulated to be under selection.

**Figure 9.**
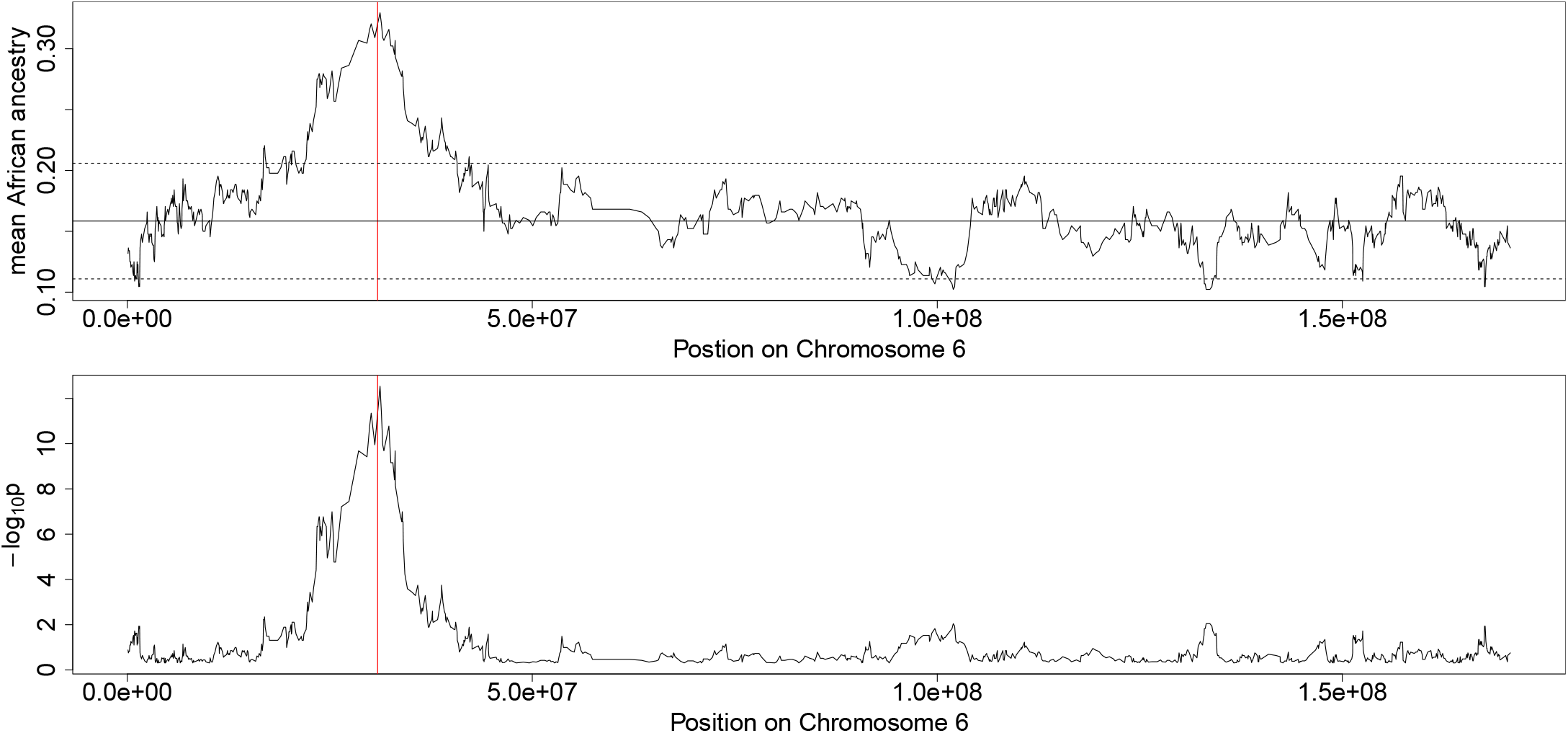
Mean Ancestry in a Wright-Fisher simulation on Chromosome 6 with positive selection at a single locus. There is a high and wide spike at the locus under selection. Note the width of the resulting spike due to hitchhiking of neighbouring loci. The solid vertical line is the mean ancestry outside the selected region and the dashed lines denote ± 2 standard deviations.

Importantly, when we then set real haplotypic data along the so-simulated ancestry segments but with *s* = 0 (no postadmixture selection effect) using donors from Southern Europe and sub-Saharan Africa and then run MOSAIC on this simulated data we do not infer a spike of African ancestry at the HLA.

## Footnotes

1 The limit to how many sources we model is a computational one as we necessarily must sum over all possible ancestry pairs for each forward / backward pass of the algorithm at each gridpoint; thus the overhead scales as 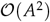 where *A* is the number of ancestries we model.

2 More correctly we could sample ancestries along the genome. However, using maximal ancestry assignment will perform a similar averaging along the genome and is far faster.

3 In practice we only assign local ancestries when the probability of assignment is greater than 0.8.

